# IF3 licenses newly made 30S subunits for translation during stress

**DOI:** 10.1101/655696

**Authors:** Indra Mani Sharma, Sarah A. Woodson

## Abstract

Bacterial ribosome biogenesis and translation occur in the same cellular compartment. Therefore, a biochemical gate-keeping step is required to prevent immature ribosomes from engaging in protein synthesis. Here, we show that the abundant ribosome assembly factor, RbfA, creates this checkpoint by suppressing protein synthesis by immature *E. coli* 30S subunits. After 30S maturation, RbfA is released by initiation factor 3 (IF3), which remains bound to 30S subunits to promote translation initiation. Genetic interactions between RbfA and IF3 show that IF3 is important for RbfA release during logarithmic growth. Moreover, IF3 is the main pathway for RbfA release in stationary phase when the activity of a less abundant RbfA-release factor, RsgA GTPase, is inhibited by the alarmone (p)ppGpp. By gating the transition from 30S biogenesis to translation initiation, RbfA and IF3 maintain the integrity of bacterial protein synthesis under a range of growth conditions and especially under stress.

## INTRODUCTION

Unlike yeast and other eukaryotes, ribosome biogenesis in bacteria occurs in the same cell compartment as protein synthesis (Klinge and Woolford, 2019; Shajani et al., 2011). Although bacteria lack membrane compartments, they must nevertheless encode robust biochemical checkpoints that gate the transition from ribosome biogenesis to translation. A barrier that prevents immature ribosomes from entering protein synthesis is essential for all cells, because translation by immature ribosomal subunits is inefficient and error prone (Andrade et al., 2018; ClatterbuckSoper et al., 2013; Cole et al., 2009; Davies et al., 2010; Roy-Chaudhuri et al., 2010; Soudet et al., 2010). How immature ribosomes are prevented from initiating translation in bacteria is not well understood. It is also not known whether the same checkpoint mechanism operates during log phase growth and in poor growth conditions that cause immature subunits to accumulate (Shajani et al., 2011).

In yeast, 40S ribosome assembly factors act as fidelity checkpoints at the last stages of pre-40S maturation prior to translation initiation (Strunk et al., 2011, 2012), through the formation of 80S-like complexes. These late assembly factors interact with the regions of the pre-40S ribosome that are recognized by translation initiation factors. A similar quality control step has not been clearly demarcated in bacteria (Datta et al., 2007; Guo et al., 2011). The binding sites of bacterial assembly factors also overlap the binding sites of translation initiation factors IF1, IF2 and IF3 (Datta et al., 2007; Guo et al., 2011; Hussain et al., 2016), however, suggesting that bacterial assembly factors may also prevent translation initiation by immature subunits. It is not known which factors mark the end of 30S biogenesis and the start of translation initiation.

Ribosome binding factor A (RbfA) is a strong candidate for the last gatekeeper in 30S biogenesis. The most abundant 30S subunit assembly factor, RbfA overexpression suppressed genetic defects in pre-16S processing (Dammel and Noller, 1995; Gibbs et al., 2017; Thurlow et al., 2016) while *rbfA* deletion impaired 30S biogenesis at low temperatures (Jones and Inouye, 1996; Xia et al., 2003). RbfA was associated with late assembly intermediates (pre-30S) and with mature 30S subunits, but not with 70S ribosomes or polysomes (Connolly and Culver, 2013; Dammel and Noller, 1995; Goto et al., 2011). A low-resolution cryo-electron microscopy structure of a 30S•RbfA complex showed that RbfA binds near the mRNA decoding A- and P-site and displaces the top of 16S helix (H) 44 and H45, rendering the complex unsuitable for joining with 50S subunits (Datta et al., 2007).

The exclusion of RbfA from 70S ribosomes (Connolly and Culver, 2013; Dammel and Noller, 1995; Goto et al., 2011) indicates that RbfA must be released before 30S subunits can initiate translation. RbfA is known to be released from mature 30S subunits by the GTPase RsgA (YjeQ) (Goto et al., 2011). In current models, GTP hydrolysis induces a conformational change within RsgA that promotes the release of RbfA and RsgA. Dissociation of RbfA and RsgA allows 16S helices H44 and H45 to dock with the 30S platform, making the 30S subunit suitable for translation (Goto et al., 2011; Guo et al., 2011; Pedro Lopez-Alonso et al., 2017; Razi et al., 2017).

Despite the well-characterized activity of RsgA GTPase, several observations suggested to us that additional *E. coli* proteins displace RbfA from 30S ribosomes. First, RsgA is nonessential, and the level of RsgA is about tenfold less than the amount of RbfA during logarithmic growth (Thurlow et al., 2016). Second, it is not known what prevents RbfA from rebinding recycled 30S subunits. Additionally, the GTPase activity of RsgA that is necessary for RbfA release was shown to be inhibited by (p)ppGpp (Corrigan et al., 2016), which accumulates during stationary phase (Peterson et al., 2005; Potrykus and Cashel, 2008). These observations suggested that *E. coli* employs a second RbfA-release factor, especially under adverse conditions.

To test this possibility, we surveyed ribosome-associated proteins for their ability to displace RbfA. Among the proteins tested, IF3 was uniquely able to release RbfA from fully mature 30S subunits but not from immature pre-30S complexes. We also found that RbfA inhibits protein synthesis by pre-30S subunits in the presence of IF3, suggesting that RbfA acts as a gatekeeper to prevent premature entry of pre-30S subunits into the translation cycle. Biochemical and genetics results further showed that IF3 is essential for displacing RbfA during stationary phase. Altogether, the results demonstrate that RbfA and IF3 enforce the barrier between ribosome biogenesis and translation, creating a checkpoint that is sensitive to the quality of the 30S decoding site.

## RESULTS

### The multi-functional initiation factor IF3 releases RbfA from 30S subunits

We employed an ultracentrifugation pelleting assay and an ultrafiltration assay (Goto et al., 2011; Jeganathan et al., 2015; Shoemaker and Green, 2011) to investigate the binding and release of RbfA from *E. coli* ribosomal 30S subunits under various conditions. In order to distinguish RbfA from ribosomal proteins (r-proteins) of similar size, we fluorescently labeled RbfA with Cy5 at position 2 (Figure S1A), which is exposed in the NMR structure of RbfA (Huang et al., 2003) and is not essential for RbfA function (Figure S1B). Fluorescent labeling enabled accurate quantitation of bound and free RbfA.

In each pelleting assay or filtration assay, 30S•Cy5-RbfA complexes were incubated with translation initiation factors or mRNA, and the bound Cy5-RbfA was separated from the unbound protein by pelleting the reaction mixture through a sucrose cushion or by filtration. In accordance with previous data (Goto et al., 2011), we observed that RbfA remained bound to mature 30S subunits for tens of minutes (t_1/2_ > 30 min) (Figure S2A). This confirmed that RbfA forms long-lived complexes with free 30S subunits in the absence of other factors.

We next tested whether components of the translation initiation complex could displace RbfA. Filtration and pelleting assays showed that neither r-protein bS1, which acts during translation initiation, nor *sodB* mRNA could displace RbfA (Figure S2B, C). RbfA also remained stably bound to 30S subunits in the presence of either IF1 or IF2 (Figure 1A, lanes 8 and 9 and Figure 1C). By contrast, we observed that RbfA dissociated from 30S subunits in the presence of 4 µM IF3, as demonstrated by a >80% decrease in the Cy5 fluorescence intensity in the pelleted fraction after ultracentrifugation (Figure 1A, lane 10 and Figure 1C). The addition of IF1 or IF2 to reactions with IF3 did not affect the ability of IF3 to promote dissociation of RbfA from 30S subunits, suggesting that IF3 acts alone on the 30S•RbfA complex (Figure S2A). We confirmed that the ability of IF3 to remove RbfA depends on its association with the 30S subunit, since IF3-K110L, which does not bind 30S ribosomes (De Bellis et al., 1992), was not able to remove RbfA (Figure 1B, lane 9 and Figure 1C). To confirm that the RbfA release promoted by IF3 is meaningful, we compared this reaction to the known RbfA release by the RsgA GTPase. RsgA was more efficient at RbfA release; in the presence of 5 µM GTP, 500 nM RsgA was as effective as 4 µM IF3 (Figure 1B, lane 10 and Figure 1C). However, IF3 levels are typically ∼20 times higher than RsgA levels during logarithmic growth; relative to 30S subunits the approximate copy numbers are 100:20:8–10:1 30S:IF3:RbfA:RsgA (Gibbs et al., 2017; Howe and Hershey, 1983; Liveris et al., 1991; Thurlow et al., 2016). Thus, these experiments suggested that IF3 provides an equally likely mechanism for removing RbfA from 30S subunits.

**Figure 1.**
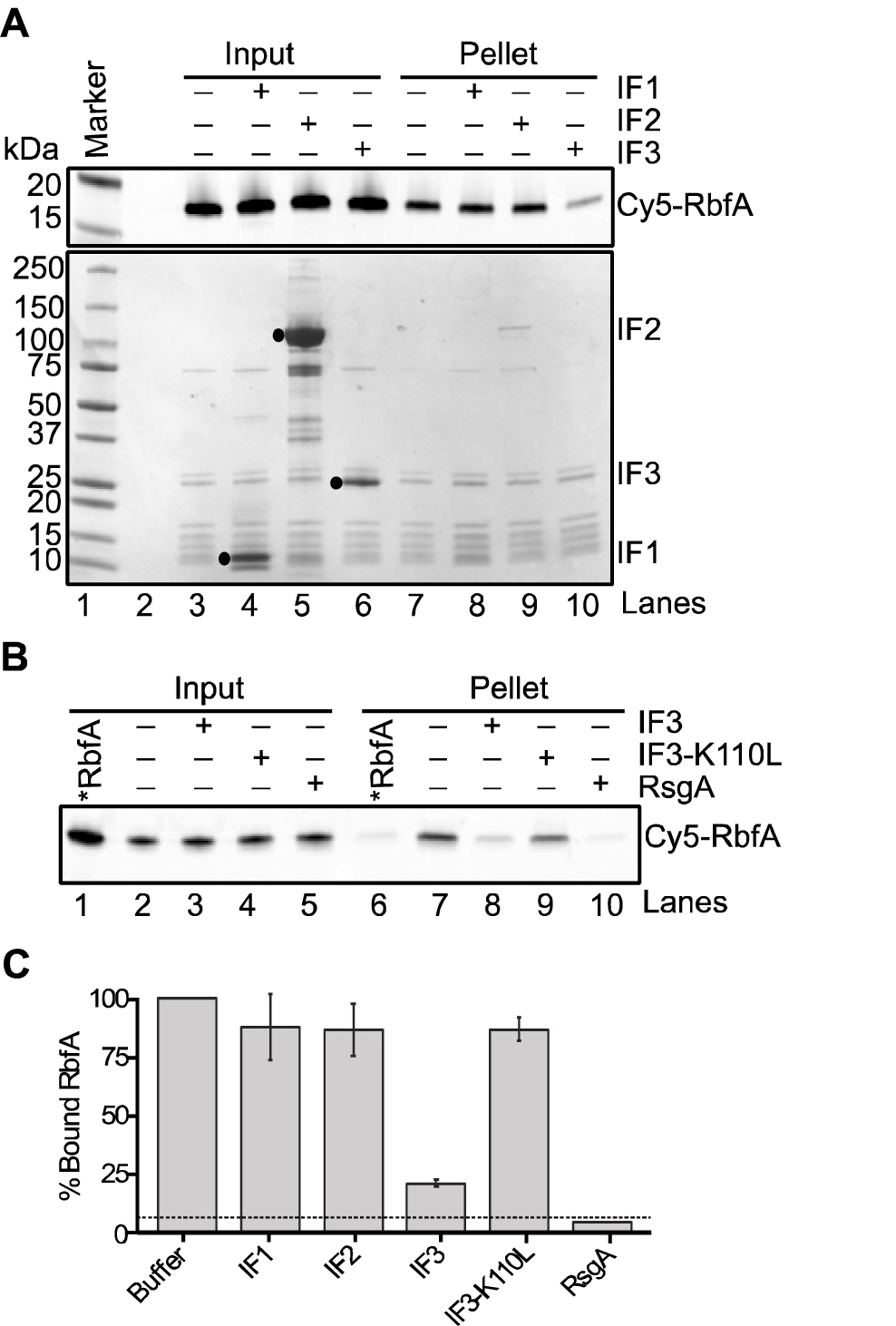
Initiation factor 3 releases RbfA from 30S subunits. (A) Results of pelleting assay showing the release of Cy5-RbfA from 30S•Cy5-RbfA complexes in the presence of initiation factors. Only IF3 was able to displace of RbfA from 30S subunits (lane 10). A complex of mature 30S (200 nM) and Cy5-RbfA (400 nM) was formed and unbound Cy5-RbfA was removed by filtration (input) before complexes were challenged with initiation factors (4 µM). Proteins were resolved by 4-20% SDS PAGE. Top panel, Cy5-RbfA fluorescence; bottom panel, Coomassie stain. Initiation factors (input) are indicated with black dots. (B) Pelleting assay showing that a non-binding IF3 mutant (IF3-K110L) cannot release of RbfA from 30S subunits (lane 9). RsgA (500 nM) and GTP (5 µM) was used as a positive control (lane 10). *RbfA (lane 1 & 6); Cy5-RbfA without 30S subunits. (C) Fraction of bound Cy5-RbfA remaining, from (A) and (B). Mean and s.d., *n* = 3 independent trials. Dotted line indicates ∼5% Cy5-RbfA background in reactions lacking 30S subunits.

### Conformational change of full length IF3 is required for RbfA release

IF3 is a 180 amino acid protein with two globular domains separated by a flexible linker. The C-terminal domain, which performs many of the functions of full-length IF3 including selection of the correct start codon (Petrelli et al., 2001), can bind the 30S subunit alone. The isolated N-terminal domain does not bind 30S subunits, but is nonetheless required for wild-type growth (Ayyub et al., 2017). Upon binding to 30S subunits, the N- and C-terminal domains of IF3 can adopt extended, intermediate, and compact conformations, owing to the dynamics of the inter-domain linker that are essential for IF3′s function in translation initiation (Elvekrog and Gonzalez, 2013).

Given these properties of IF3, we asked whether the conformational dynamics of IF3 is needed to promote the release of RbfA from 30S subunits. We used the IF3 mutation Y75N, which is located in the highly conserved linker region and is known to be critical for the start codon selection activity of IF3 (Maar et al., 2008; Sussman et al., 1996). This mutation does not affect IF3 binding to 30S subunits, but it disfavors the extended conformation of IF3 (Elvekrog and Gonzalez, 2013). We found that the Y75N mutation reduced the amount of RbfA released compared to WT IF3 (Figure 2A and 2C), suggesting that IF3′s ability to fluctuate into the extended conformation is required for optimal release of RbfA from 30S subunits. To completely abrogate the conformational dynamics, we separated IF3 into N-terminal and C-terminal fragments and tested their ability to remove RbfA from 30S subunits (Figure 2B and 2C). As expected, the separated domains individually or in combination failed to remove RbfA from 30S subunits, suggesting that conformational changes of the full-length IF3 are important for displacing RbfA.

**Figure 2.**
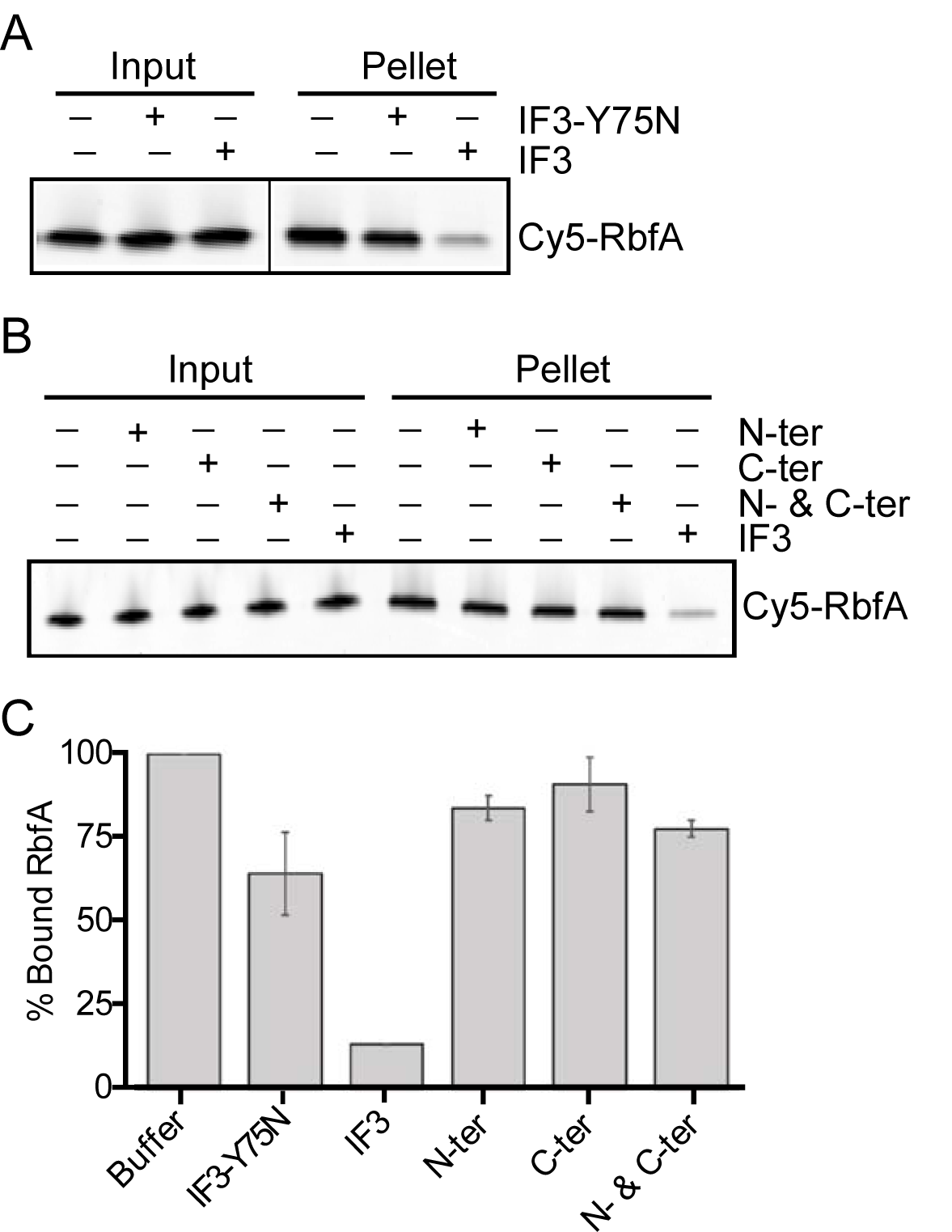
Conformational change in full-length IF3 is required for RbfA release. (A) Results of pelleting assay showing that a mutation in the linker region of IF3 (IF3-Y75N) reduces Cy5-RbfA release. Experiment performed as in Fig. 1. (B) Separated N- and C-terminal domains of IF3 alone or in combination cannot promote RbfA release. (C) Fraction of bound Cy5-RbfA in (A) and (B), as in Fig. 1. Error bars, ±s.d. *n* = 3.

### Timing the release of RbfA during 30S subunit biogenesis

To identify at which step RbfA acts during 30S biogenesis, we sought to determine the influence of 30S subunit composition on the process of RbfA release by IF3. We found that RbfA does not bind with naked 16S 5’ domain RNA, 16S or 17S rRNA, nor with early 30S assembly intermediates reconstituted *in vitro* (Figure S3A, B, and C), consistent with previous studies showing that RbfA acts during late 30S subunit assembly or maturation (Connolly and Culver, 2013; Dammel and Noller, 1995; Datta et al., 2007; Goto et al., 2011).

Late 30S assembly intermediates were purified from a *ΔrbfA* strain grown at low temperature that causes pre-30S complexes to accumulate (ClatterbuckSoper et al., 2013). These pre-30S particles contain unprocessed 17S pre-rRNA and lack some r-proteins (Figure 3A). Cy5-RbfA was allowed to bind pre-30S particles as before, then challenged with IF3, followed by ultracentrifugation. Although RbfA readily binds pre-30S particles, IF3 was not able to promote the release of RbfA from these complexes. Thus, IF3 preferentially releases RbfA from mature 30S subunits (Figure 3B), as shown previously for RsgA (Goto et al., 2011).

**Figure 3.**
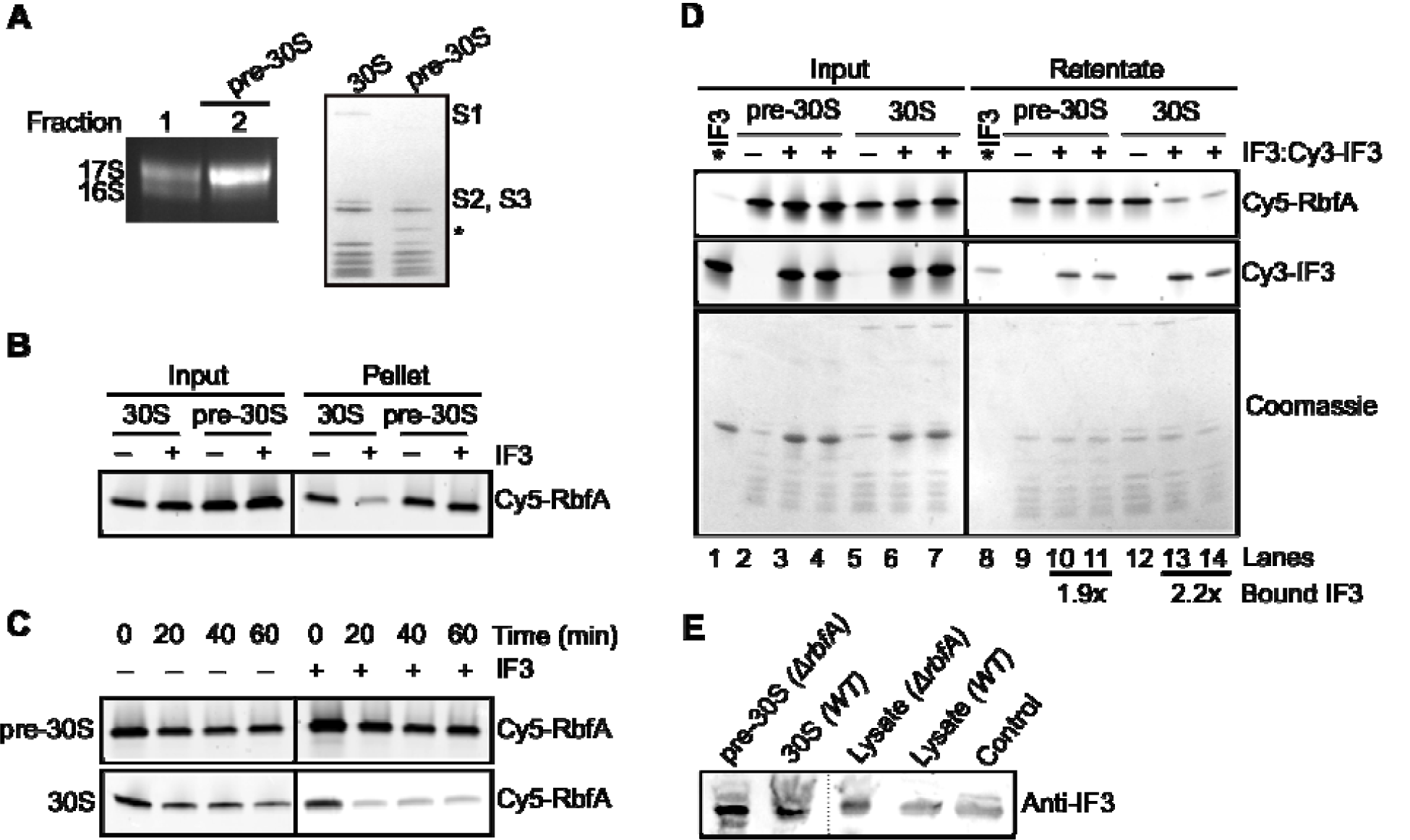
IF3 cannot release RbfA from immature pre-30S subunits. (A) A 1.5% agarose-TAE gel showing the rRNA composition of pre-30S _*ΔrbfA*_ particles (left panel). The fraction 2 containing more than 90% 17S pre-rRNA (pre-30S) was used for further assays. 4-20% SDS PAGE comparing the protein composition of mature 30S and pre-30S _*ΔrbfA*_ particles (right panel). The absence of S1, S2, and S3 is evident in the pre-30S particles. The identity of the extra band (*) in the pre-30S fraction is not known. (B) Pelleting assay; Cy5-RbfA (400 nM) was complexed with mature 30S or pre-30S subunits (200 nM) and challenged with IF3 (4 µM) as in Fig.1. (C) Kinetics of RbfA dissociation from pre-30S or 30S subunits in the absence or presence of IF3. After Cy5-RbfA•30S complexes were formed and excess RbfA removed by filtration (0 min lanes), they were incubated an additional 20 – 60 min at 37 °C, with or without 4 µM IF3, before filtration a second time to determine the amount of Cy5-RbfA that remained bound. The 0 min lanes that were filtered only once contain about 30% more RbfA than the remaining samples, which were filtered twice. (D) Filtration assay as above with Cy5-RbfA and pre-30S or 30S subunits. Top panel, Cy5-RbfA in the retentate; middle panel, Cy3-IF3 in the retentate; lower panel, Coomassie stain. *IF3 (lane 1 and 8); Cy3-IF3 without 30S subunits and Cy5-RbfA. Average fold change in bound IF3 (over *IF3 background in lane 8) was 1.9 ± 0.15 in lanes 10 and 11 and 2.2 ± 0.89 in lanes 13 and 14. IF3 was added as a mixture of unlabeled IF3 (80%) and Cy3-IF3 (20%). Cy5-RbfA was scanned with 600 PMT, whereas Cy3-IF3 (Input) with 400 PMT and Cy3-IF3 (retentate) with 500 PMT. (E) Anti-IF3 Western blot showing the presence of IF3 in the pre-30S and 30S fractions from BX41 (Δ*rbfA*) and BW25113 (WT) cells (left). Right, Δ*rbfA* and WT lysates; control, purified IF3.

To understand why RbfA could not be released from pre-30S subunits, we compared the lifetimes of RbfA bound to pre-30S and mature 30S subunits by removing free RbfA by ultrafiltration at various times. Figure 3C shows that the half-life of the pre-30S•RbfA complex was > 60 min as observed previously (Goto et al., 2011), and was unaffected by the presence of IF3. Whereas, the amount of RbfA bound to mature 30S subunits significantly decreased after 20 min in the presence of IF3. These experiments suggested that the structure of the pre-30S particles either stabilizes RbfA binding so that it cannot be removed by IF3, or prevents IF3 from acting on the complex.

To test whether IF3 fails to release RbfA because IF3 cannot bind pre-30S complexes, we monitored the release of Cy5-RbfA in the presence of Cy3-IF3, followed by detection of Cy3 signal to determine whether IF3 was retained with the 30S or pre-30S complexes after filtration. Both complexes were able to bind IF3 in the presence of RbfA (Figure 3D). This result suggested that either conformational changes in the linker of IF3 are compromised on pre-30S subunits, or that these conformational changes occur but cannot dislodge RbfA from pre-30S subunits (Figure 3D, Lanes 10 and 11). Importantly, in a similar experiment with mature 30S subunits, we observed that IF3 remains bound to 30S subunits after removing RbfA (Figure 3D, lanes 13 and 14). Thus, IF3 can bind both mature and immature complexes, but cannot displace RbfA from pre-30S particles.

We sought to determine if IF3 binds pre-30S particles *in vivo* by analyzing purified pre-30S or 30S samples for the presence of IF3 by Western blotting (Figure 3E). The complexes were fractionated by sucrose gradient sedimentation and validated by measuring rRNA and protein composition (as in Figure 3A). We found that IF3 co-purifies with pre-30S subunits in an *ΔrbfA* strain. Additionally, IF3 was previously detected by Western blotting in pre-30S particles in a *ΔybeY* strain (Davies et al., 2010) and by mass spectrometry in pre-30S particles containing 16S mutations that block folding of a helix junction (Sharma et al., 2018) (Figure S4). Together, these data confirmed that the interaction of IF3 with pre-30S particles is physiologically measurable. Furthermore, pre-30S subunits have been found in 70S and polysome fractions of *ΔrbfA, ΔrimM, ΔybeY*, and *ΔrpsO* strains (ClatterbuckSoper et al., 2013; Davies et al., 2010; Roy-Chaudhuri et al., 2010), suggesting that some pre-30S complexes are competent for translation and therefore must interact with IF3.

### RbfA release marks the transition from ribosome biogenesis to translation initiation

Methylation of 16S A1518 and A1519 in H45 by the conserved RNA methylase KsgA is thought to be one of the last steps in the 30S subunit biogenesis preceding translation initiation (Connolly and Culver, 2013; Connolly et al., 2008; Thammana and Held, 1974). Therefore, we wondered if H45 methylation promotes RbfA release by IF3. First, we used primer extension to confirm that pre-30S complexes purified from an Δ*rbfA* strain are methylated by KsgA, as indicated by the presence of a reverse transcription stop at residue A1519 (Figure 4A, lane 4). This showed that RbfA is not required for H45 methylation, consistent with the proposal that RbfA acts downstream of KsgA (Connolly and Culver, 2013).

**Figure 4.**
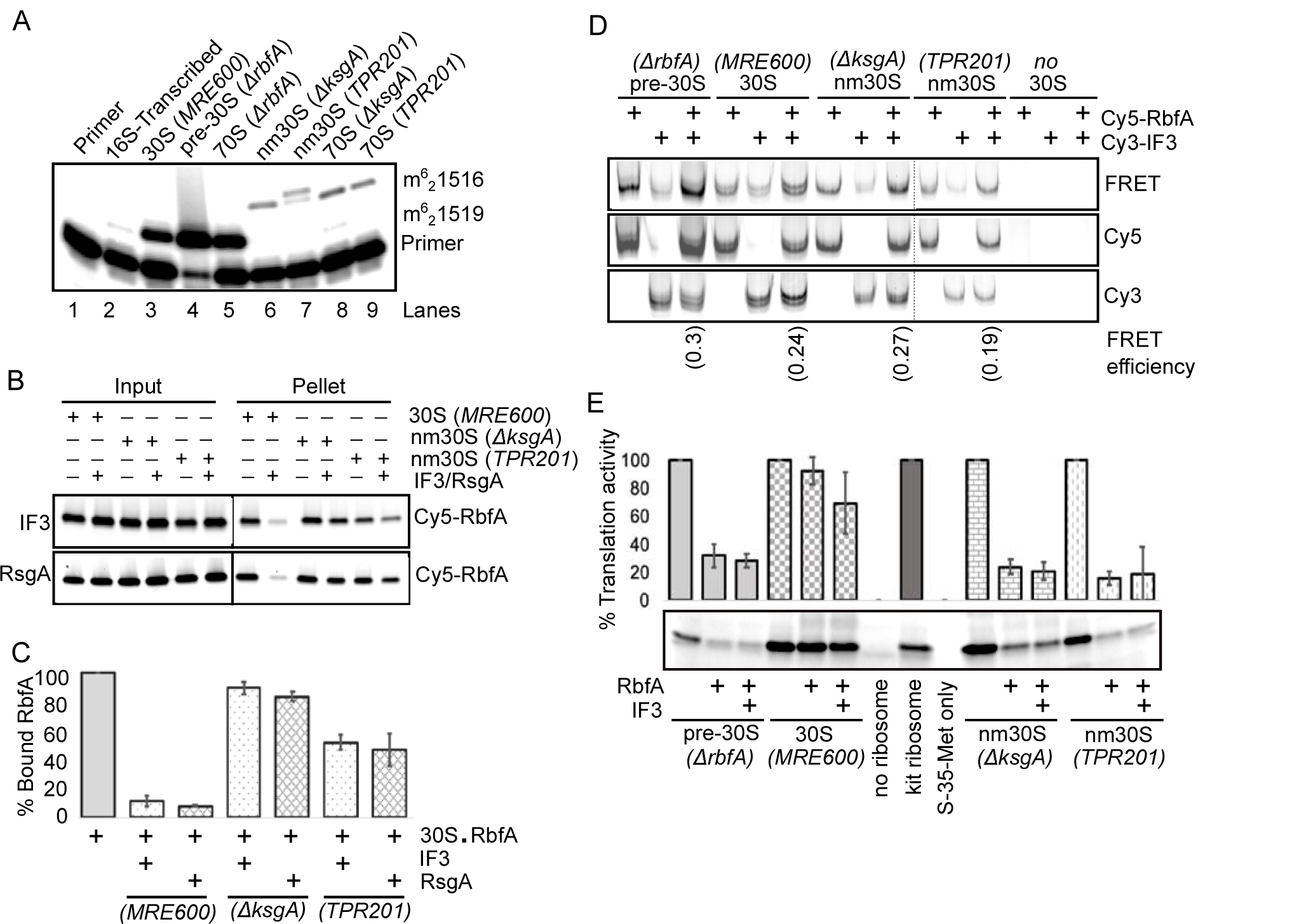
RbfA inhibits translation by immature 30S ribosomes. (A) Methylation of 16S A1519 in pre-30S and 30S subunits was measured by primer extension (see Methods). Pre-30S subunits from BX41 (*ΔrbfA*) were methylated by KsgA (lane 4); nm30S subunits from ΔksgA (JW0050-3) and TPR201 bearing an inactive *ksg*A allele were unmethylated (lanes 6 and 7). (B) Pelleting assay showing that both IF3 and RsgA failed to promote release of Cy5-RbfA from unmethylated *ΔksgA* nm30S subunits and poor release from non-methylated TPR201 nm30S subunits. (C) % Bound RbfA relative to 30S control from (B); mean and s.d.; *n* = 2. (D) Native PAGE colocalization showing that Cy5-RbfA and Cy3-IF3 can bind pre-30S or 30S subunits simultaneously. 30S complexes were purified from the strains shown and incubated with Cy5-RbfA and Cy3-IF3 before native PAGE. Panels show Cy3 intensity upon Cy3 excitation (bottom), Cy5 intensity upon Cy5 excitation (middle), and FRET to Cy5 upon Cy3 excitation (top). The FRET efficiencies are indicated at the bottom. (E) *In vitro* translation by pre-30S (methylated), nm30S (unmethylated) and mature, methylated 30S subunits in the presence of RbfA or RbfA plus IF3. Top, average and s.d. of two experiments. Bottom, amount of ^35^S-labeled DHFR protein produced. Immature or unmethylated subunits are active but are inhibited by RbfA, which remains bound. However, 30S (methylated) subunits, from which RbfA was removed by IF3, were unaffected by RbfA.

Next, to determine whether H45 methylation is required for RbfA release, we purified unmethylated <u>near-mature</u> 30S (nm30S) subunits from two strains lacking KsgA activity: *ΔksgA* deletion (Baba et al., 2006) and TPR201, which harbors a catalytically dead *ksgA* allele (Andrésson and Davies, 1980). nm30S subunits purified from these strains were not methylated at A1519 (Figure 4A, lanes 6 and 7), although they contain the full complement of r-proteins (Connolly and Culver, 2013) and the mature 16S rRNA 5′ end (Figure S5A). Pelleting assays showed that RbfA was able to bind to these nm30S particles, but its release by IF3 was compromised. IF3 was unable to promote release of RbfA from nm30S subunits from the *ΔksgA* strain, and only partially able to release RbfA from TPR201 nm30S subunits (Figure 4B, upper panel and Figure 4C). The absence of H45 methylation also reduced the ability of RsgA to release RbfA from both types of nm30S subunits (Figure 4B, lower panel and Figure 4C). Thus, while methylation by KsgA is not essential for RbfA recruitment, it is important for RbfA release.

We next examined the difference in IF3 binding to nm30S subunits to determine if this could explain IF3’s reduced ability to promote the release of RbfA from these subunits. nm30S subunits from *ΔksgA* and TPR201 strains were incubated with Cy5-RbfA and Cy3-IF3 and then subjected to native PAGE (Figure 4D). We observed colocalization of Cy3 and Cy5 fluorescence in the native gel for all of the 30S complexes tested, indicating that IF3 and RbfA can both bind to methylated and non-methylated 30S subunits. Furthermore, we observed FRET from Cy3-IF3 to Cy5-RbfA (*E*_FRET_ ∼ 0.2 to 0.3), indicating that IF3 and RbfA can bind 30S, nm30S or pre-30S complexes at the same time. IF3 interacted less tightly with unmethylated nm30S subunits than with 30S_WT_ subunits, however, judging from the intensities of individual protein bands. Connolly & Culver (2013) also showed that IF3 binds less tightly to nm30S subunits from Δ*ksgA* cells, supporting our observation. Together, these data show that KsgA methylation of H45 enhances IF3 binding, and that methylation is important for the efficient removal of RbfA from 30S subunits. These results explain why RbfA overexpression is toxic to cells lacking KsgA (Connolly and Culver, 2013), because RbfA cannot be released from nm30S subunits.

### RbfA suppresses translation by pre-30S ribosomes

We previously observed that pre-30S ribosomes are partially active in protein synthesis and enter the polysome fraction in an *E. coli* strain lacking RbfA (ClatterbuckSoper et al., 2013), supporting the idea that RbfA normally prevents translation by immature subunits. Since IF3 and RbfA can bind 30S subunits simultaneously, we wondered if RbfA release is required for efficient translation initiation. To test this, we compared the activity of pre-30S (*ΔrbfA*), nm30S (*ΔksgA* and TPR201) and mature 30S subunits in an *in vitro* translation assay using purified components (Shimizu et al., 2001) in the presence or absence of excess RbfA (Figure 4E). Pre-30S_*ΔrbfA*_ subunits were intrinsically less active (29 ± 0.3%) than mature 30S subunits, consistent with previous work (ClatterbuckSoper et al., 2013). In contrast, nm30S subunits had normal activity. Importantly, the addition of RbfA substantially inhibited translation by pre-30S_*ΔrbfA*_ and nm30S (*ΔksgA* and TPR201) complexes, while it did not inhibit translation by WT 30S_*MRE600*_ subunits (Figure 4E). Increasing the concentration of IF3 in the translation assay did not improve the translation efficiency for any of the 30S complexes tested. We concluded that IF3’s inability to release RbfA from immature particles (pre-30S_*ΔrbfA*_, nm30S_*ΔksgA*_, and nm30S_*TPR201*_), even when IF3 was provided in excess, markedly decreased the activity of these subunits. These data suggest that RbfA senses incomplete maturation of 30S subunits (as in case of pre-30SΔ_*rbfA*_ and nm30S from *ΔksgA* and TPR201 strains) and directly prevents their premature entry into translation, providing a last checkpoint for ribosome biogenesis.

### RbfA and IF3 are genetically linked

The results above show that IF3 can displace RbfA from mature 30S subunits *in vitro*, but not from immature subunits. To determine if this activity of IF3 is important *in vivo*, we looked for a genetic interaction between RbfA and IF3. Since IF3 is essential, we used a non-lethal IF3-Y75N mutation (*infC*362; (Sussman et al., 1996)), which is fortuitously defective in its ability to release RbfA from 30S subunits (Fig. 2A). Figure 5A shows that the *infC362* strain grows similarly to the parental *E. coli* strain. When these strains were transformed with a plasmid expressing RbfA (p15BHA), however, leaky expression of RbfA in the absence of IPTG inhibited the growth of cells expressing IF3-Y75N but not cells expressing wild type IF3 (Figure 5A). This level of RbfA expression is enough to complement an *E. coli* Δ*rbfA* strain (Dammel and Noller, 1995), suggesting that RbfA and IF3 are genetically linked. The leaky expression of RbfA was also toxic when the *infC*362 cells were grown in minimal media (Figure 5B), which slows ribosome biogenesis and makes cells more dependent on ribosome assembly factors. The genetic interaction between IF3 and RbfA was confirmed by plate assays, which also showed that RbfA over-expression induces the cold sensitivity of *infC*362 cells (Figure 5C). We concluded that wild type IF3 plays an important role in removing RbfA from 30S subunits, which becomes even more important at low temperature or when nutrients are limited.

**Figure 5.**
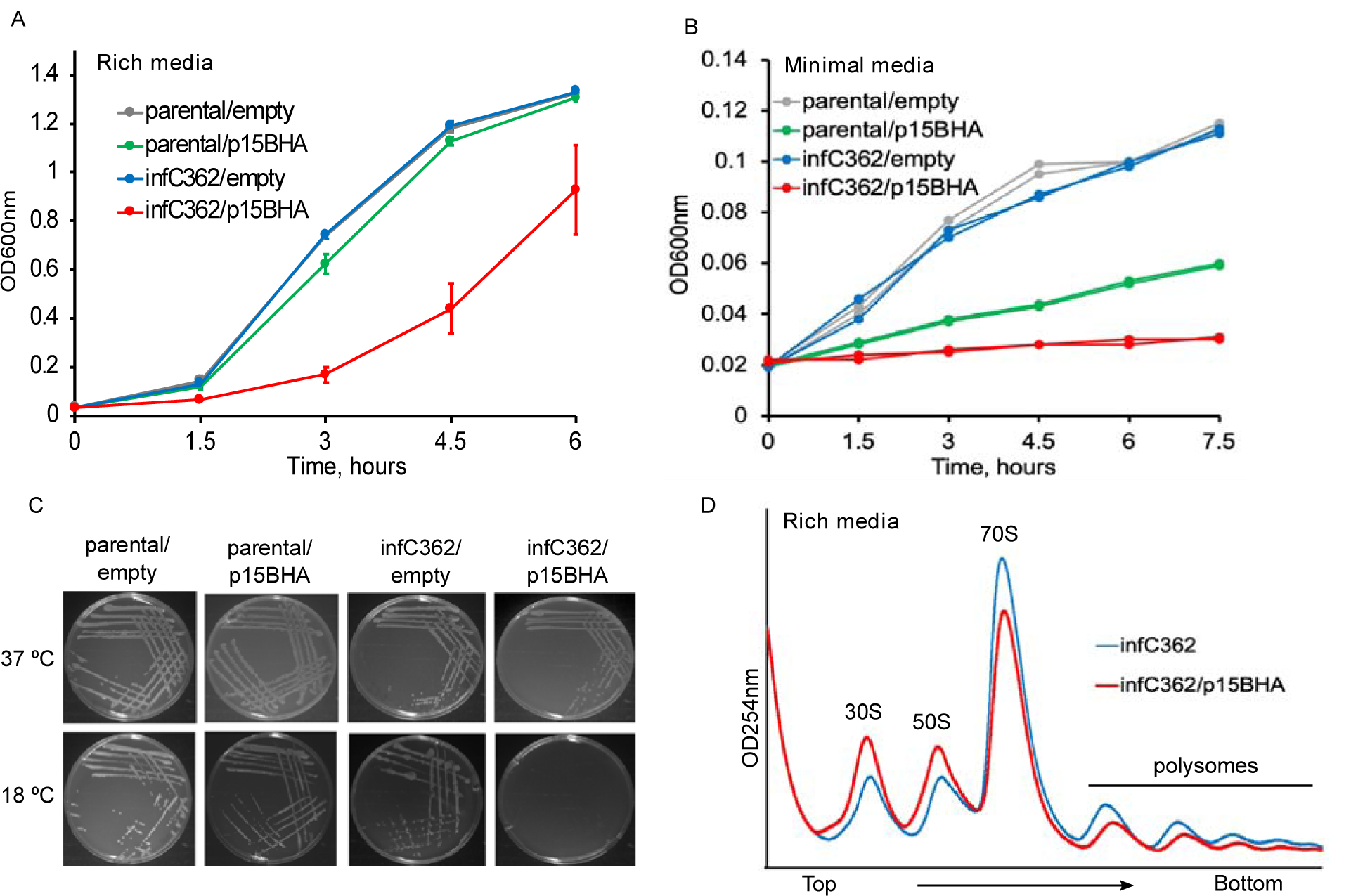
Genetic interaction between RbfA and IF3. (A) Growth curves of *E. coli* strains parental (JK382)/empty vector (pSE420), infC362 (JK378)/empty vector, and parental and infC362 transformed with p15BHA (for over-expression of RbfA-HA) in rich LB media at 37 °C. Growth was monitored by recording OD_600nm_. Error bars, s.d. of four replicates. (B) Growth curves of strains as shown in (A) in minimal MOPS media (pH 7.2) supplemented with 0.4% glucose at 37 °C; *n* = 2. (C) Growth of strains as shown in (A) on LB agar media plates at 37 °C and 18 °C; *n* = 3. (D) Polysome profiles of strains infC362 and infC362/p15BHA, which were grown under similar conditions as shown in (A) and collected at OD_600nm_ ∼0.2. Experiment was performed in duplicate.

To pinpoint the reason for RbfA toxicity in the IF3-Y75N background, the polysome profile of *infC*362/p15BHA strain was compared with *infC362* strain. Figure 5D shows that excess RbfA reduced the size of the 70S and polysome peaks, with a concomitant increase in the amounts of free 30S and 50S ribosome subunits. This is consistent with an inability of IF3-Y75N to displace RbfA, which in turn stabilizes free 30S subunits. This reduction in the numbers of active ribosomes was not due to a defect in 30S biogenesis because we did not observe a defect in 16S rRNA 5′ processing as judged by primer extension (Figure S5B).

### IF3 is the predominant pathway for RbfA release during stationary phase

Since the genetic interaction of IF3 and RbfA is stronger in low nutrient conditions, we reasoned that IF3 may be the dominant factor that releases RbfA during stationary phase, when the GTPase activity of RsgA is inhibited by (p)ppGpp (Corrigan et al., 2016). To test this possibility, we measured RbfA release in a mixture containing combinations of RsgA, GTP, ppGpp and IF3. We found that ppGpp inhibits the release activity of RsgA by ∼55% (Figure 6A, lane 10 and Figure 6B), yet did not inhibit IF3’s ability to release RbfA from 30S subunits (Figure 6A, lane 11 and 12 and Figure 6B). These data suggest that IF3 can promote the release of RbfA when ppGpp levels are high enough to inhibit RsgA, as is the case during stationary phase.

**Figure 6.**
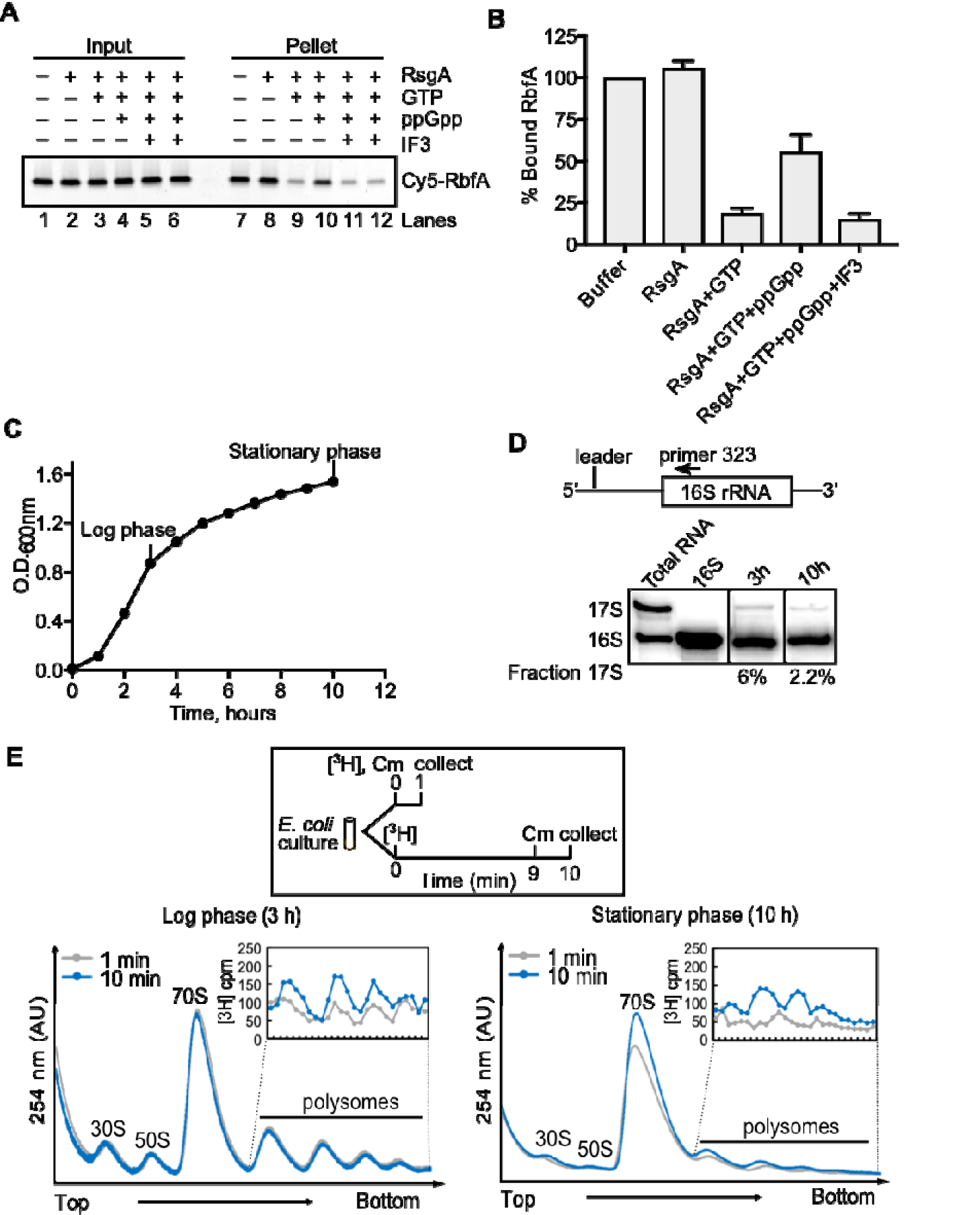
IF3 is the predominant pathway during stationary phase. (A) Pelleting assay showing that RbfA release by 200 nM RsgA (with 50 µM GTP) is inhibited by 1.5 mM ppGpp (lane 10). 5 µM IF3 can release of RbfA from 30S subunits under this condition (lane 11 and 12). (B) Quantification of experiments performed in (A). Error bars are the s.d. of the average from two experiments. (C) Growth of WT *E. coli* (BW25113) cells in liquid LB medium. (D) Primer extension on total RNA purified from cells grown in (C) at 3h and 10h to show the presence of unprocessed 17S pre-rRNA as a proxy of ribosome biogenesis. (E) A comparison of polysome profiles during log phase (left panel) and stationary phase (right panel) from tritium labeled cells. Cells were grown 3 h or 10 h, and collected at 1 and 10 min after the addition of ^3^H-uridine and 1 min after treatment with chloramphenicol. Inset: incorporation of ^3^H-uridine into polysomes.

Because ribosome biogenesis is sharply reduced in stationary phase or when nutrients are limiting (Dennis and Bremer, 2008; Gourse et al., 1996; Murray et al., 2003), we asked whether a checkpoint for biogenesis is needed under these conditions. To measure the amount of ribosome biogenesis during stationary phase, we purified total rRNA from cells in mid-log or stationary phase (Figure 6C) and measured the amount of unprocessed 17S pre-rRNA by primer extension (ClatterbuckSoper et al., 2013; Gupta and Culver, 2014). Immature 17S rRNA was present during both growth stages tested, although the proportion of immature rRNA was three times lower during stationary phase (2% vs. 6%; Figure 6D). We next tracked *de novo* ribosome synthesis by pulse labeling the rRNA with ^3^H-uridine in cells in log phase (3 h, OD_600nm_ ∼0.6-0.8) and stationary phase (10 h, OD_600nm_ ∼1.6-1.8), to determine whether newly synthesized rRNA is assembled into mature ribosomes (Figure 6E). The numbers of ribosomes and polysomes are expected to be lower in stationary phase (Reeve et al., 1984). Nevertheless, after 10 min, newly formed tritiated ribosomal subunits were observed in the polysome fractions from cells in stationary phase, indicating that ribosome biogenesis continues even when nutrients are limiting (Figure 6E).

## DISCUSSION

Here, we show that a translation initiation factor, IF3, robustly displaces RbfA, the most abundant 30S subunit assembly factor in *E. coli*. Because RbfA is only released from 30S subunits when assembly and maturation are complete, this suggests that the opposing effects of RbfA and IF3 on pre-30S particles set up a checkpoint between 30S biogenesis and protein synthesis (Figure 7). Each protein interacts with 16S H44 (decoding center), explaining how this checkpoint senses the assembly status of the 30S active site (Figure S6). After 30S maturation, a hand-off from RbfA to IF3 (30S•RbfA→30S•IF3) leads to formation of a 30S translation initiation complex.

**Figure 7.**
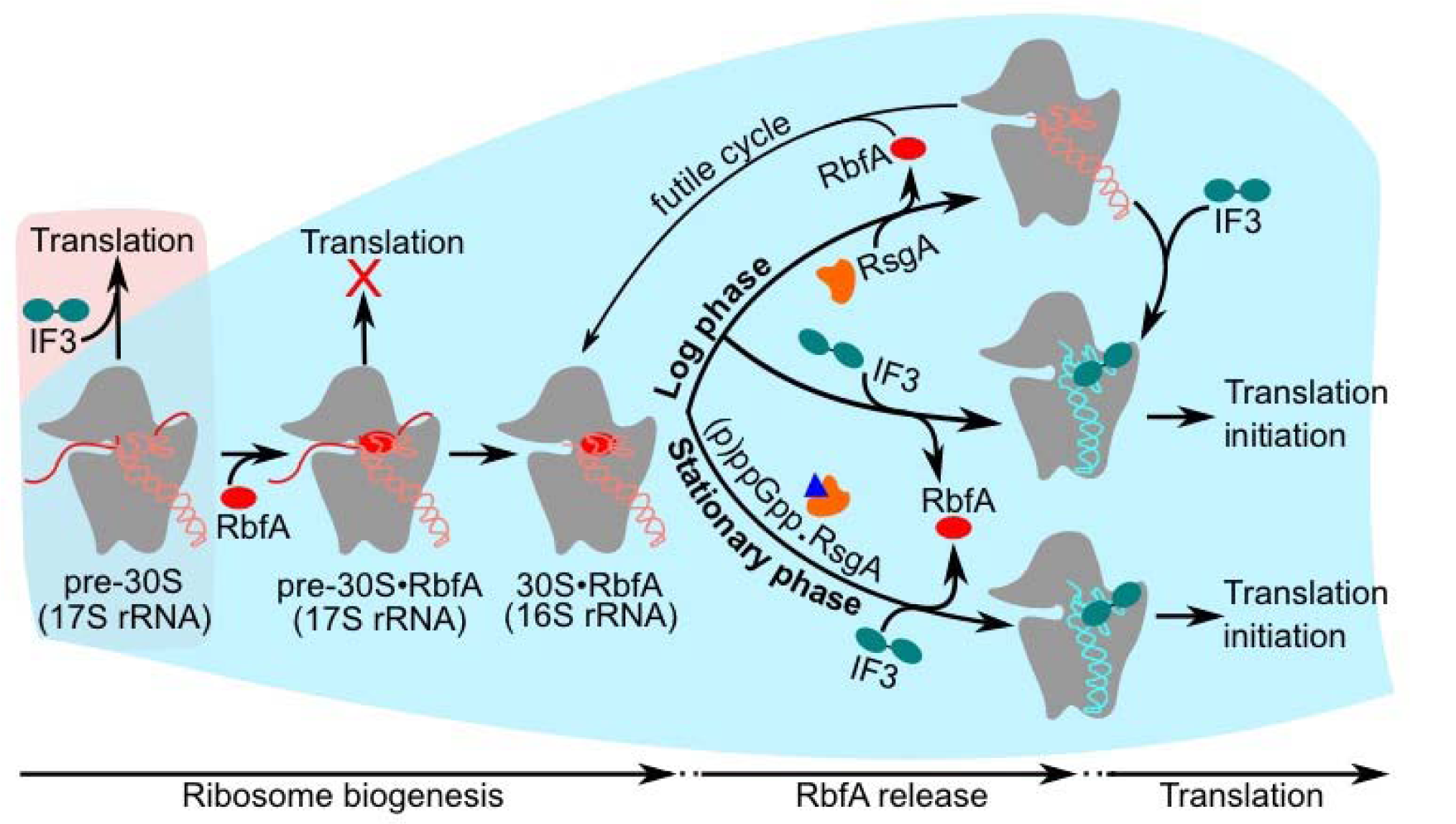
A model depicting pathways involving release of RbfA from 30S subunits. Under normal growth conditions (blue background), RbfA strongly interacts with immature pre-30S particles and prevents their entry into translation. Both RsgA and IF3 can promote the release of RbfA from mature 30S subunits. However, when RsgA releases RbfA, free 30S subunits can again bind to RbfA, leading to a futile cycle. 16S H44 and H45 in free 30S subunits can fluctuate into an undocked state (ClatterbuckSoper et al., 2013; McGinnis et al., 2015). By contrast, when IF3 releases RbfA from mature 30S subunits, IF3 remains bound and ready to form a translation initiation complex (30SIC). During stationary phase, RsgA is inhibited by (p)ppGpp and RbfA is mainly released by IF3. Under stress (red background), when the pre-30S level exceeds the amount of RbfA, some pre-30S complexes can also translate mRNAs.

### RbfA as a gatekeeper for 30S ribosomes

During active growth conditions, bacteria must ensure the fidelity of translation by preventing the entry of pre-30S particles into translation, which are inefficient and error-prone (Andrade et al., 2018; ClatterbuckSoper et al., 2013; Davies et al., 2010; Roy-Chaudhuri et al., 2010). We found that RbfA directly inhibits translation by immature pre-30S particles (Figure 4E), providing a gatekeeping mechanism. Congruent with this role, RbfA forms stable complexes with pre-30S subunits (Figure 3B, C) that are not dissociated by either RsgA or IF3 (Figure 4B, C). In the cryo-EM structure of the RbfA•30S complex, RbfA dislocates 16S helices H45 and the top of H44 (Datta et al., 2007), preventing initiator tRNA binding and the formation of bridges B2a and B3 with the 50S subunit. One possibility is that RbfA senses incomplete maturation of 30S subunits by testing the stability of interactions around the 30S decoding center, which is the last region of the 30S subunit to fold. This may also explain why RbfA release is sensitive to KsgA methylation of A1518 and A1519, which stabilizes tertiary interactions between H45 and H44 (Boehringer et al., 2012; Pedro Lopez-Alonso et al., 2017). In yeast, late assembly factors were also suggested to prevent the premature entry of pre-40S into translation cycle by blocking formation of the 43S pre-initiation complex (Strunk et al., 2011).

### Mechanism of RbfA release by IF3

It has been known for some time that RbfA does not bind with 70S ribosomes or polysomes, indicating that RbfA is released from 30S subunits before they enter translation (Connolly and Culver, 2013; Dammel and Noller, 1995; Goto et al., 2011; Jones and Inouye, 1996). Our observation that RbfA does not interact with the 30S pre-initiation complex (30SIC) (Figure S3D, lane 3) further suggested that RbfA is released before or during initiation of translation. Although IF2 was previously reported to genetically interact with RbfA (Jones and Inouye, 1996), in our assays, only IF3 is able to release RbfA from mature 30S subunits (Figure 1 and S2).

Although our results don’t reveal how IF3 displaces RbfA from mature 30S ribosomes, cryo-EM and single molecule FRET findings suggest a plausible pathway for RbfA displacement (Figure S6). Cryo-EM structures showed that IF3 initially interacts with the 30S subunit through its N-terminal domain which binds the platform near uS11 (Elvekrog and Gonzalez, 2013; Hussain et al., 2016). This interaction anchors IF3 to 30S subunits and allows for a conformational change within the linker region that extends the C-terminal domain towards the top of H44, where it reinforces docking of H44 and H45 and stabilizes the tRNA anti-codon in the P-site (Elvekrog and Gonzalez, 2013; Hussain et al., 2016; López-Alonso et al., 2017). Together these observations suggest that IF3 and RbfA exert opposite and mutually exclusive effects on docking of H44 and H45, which can explain how IF3 binding favors RbfA dissociation. In-cell footprinting indicated that H44 is at least partially unfolded in free 30S subunits (McGinnis et al., 2015) and completely undocked in pre-30S complexes (ClatterbuckSoper et al., 2013), supporting the notion that this region of the 30S ribosome can fluctuate between docked and undocked conformations.

By contrast with IF3, RsgA uses GTP hydrolysis to induce conformational changes that promote the release of RbfA (Goto et al., 2011; Guo et al., 2011; Pedro Lopez-Alonso et al., 2017; Razi et al., 2017). Although this likely requires folding and docking of 16S H44 and H45, we cannot exclude the possibility that RsgA and IF3 displace RbfA by different mechanisms. An important difference is that RsgA itself dissociates from the 30S complex upon GTP hydrolysis. By contrast, IF3 remains bound after RbfA dissociates, and could be physiologically more important for preventing RbfA rebinding and toxicity. This idea is further supported by the fact that the N-terminal region of RsgA, which is important for RbfA release, is weakly conserved or absent from many bacteria (Razi et al., 2017).

### RbfA release by IF3 increases stress tolerance

Our data indicate that RbfA release by IF3 is even more critical at low temperature and when nutrients are limited (Figure 5). Ribosome biogenesis continues during stationary phase, albeit slowly (Figure 6D and E). However, as the GTPase activity of RsgA is inhibited by rising levels of (p)ppGpp (Figure 6A and B), IF3 is increasingly needed to remove RbfA from newly made 30S subunits. RsgA has been shown to promote 70S ribosome dissociation into 30S and 50S subunits (Guo et al., 2011; Himeno et al., 2004), which may explain why inhibition of RsgA by (p)ppGpp is advantageous to the cell. During adaptation to low temperature and starvation, 70S ribosomes can also enter hibernation by forming 100S dimers (Starosta et al., 2014; Wada et al., 2000). Interestingly, IF3 is known to compete with hibernation factors for binding to ribosomes following termination, suggesting a competition between recycling and hibernation (Seshadri and Varshney, 2006; Yoshida et al., 2009). It will be interesting to see which of these pathways is more important for maintaining cell survival under stress conditions.

### Translation using pre-30S subunits is a survival strategy under stress

When pre-30S particles accumulate during stress, they participate in translation to ensure survival. The results of our in vitro translation assays support previous reports that pre-30S complexes can enter the polysome pool in the absence of RbfA or when 30S assembly has stalled (ClatterbuckSoper et al., 2013; Davies et al., 2010; Roy-Chaudhuri et al., 2010). We suggest that when the accumulation of pre-30S subunits exceeds RbfA concentrations, free pre-30S complexes which are competent can enable translation under different adverse growth conditions (Figure 7).

Together, our data support a model in which RbfA normally prevents the premature entry of pre-30S complexes into translation by exploiting the instability of 16S rRNA interactions within the immature decoding site (Figure 7). After 30S maturation, RbfA binding is destabilized, and RsgA or IF3 can release RbfA to keep up with the demand for new 30S subunits during logarithmic growth. Since IF3 remains bound with mature 30S subunits after RbfA release, the release of RbfA by IF3 therefore marks end of ribosome biogenesis and beginning of translation initiation.

## ACKNOWLDGEMENTS

The authors thank Julie Brunelle, Himani Galagali, and Laura Lessen for technical help, Drs. Gloria Culver, Ruben Gonzalez, Rachel Green, Hyouta Himeno, Isabella Möll, Harry Noller, and Umesh Varshney for the gifts of reagents and strains, Drs. Balasubrahmanyam Addepalli and Patrick Limbach for the assistance with mass spectrometry, and Drs. Allen Buskirk, Yumeng Hao, Arthur Korman, and Margaret Rodgers for helpful discussion and comments on the manuscript. This work was supported by a grant from the NIH (R01 GM60819 to SAW).

## AUTHOR CONTRIBUTIONS

I.M.S. conceived the project, performed experiments, analyzed and interpreted the results and wrote the manuscript. S.A.W. helped design the project, interpret the results and edit the manuscript.

## DECLARATION OF INTERESTS

The authors declare no competing interests.

## STAR ⍰ METHODS

Detailed methods are provided in the online version of this paper and include the following:

- KEY RESOURCES TABLE
- CONTACT FOR REAGENT AND RESOURCE SHARING
- METHODS DETAILS
  - Strains and culture conditions
  - Plate assays
  - Plasmids
  - Ribosome purification
  - Protein purification and fluorescence labeling
  - Ultracentrifugation pelleting assay and ultrafiltration assay
  - Primer extension
  - *In vitro* translation assay
  - ^3^H-Uridine pulse labeling and polysome profiling
  - Western blot
  - Native PAGE experiments

## SUPPLEMENTAL INFORMATION

Supplemental Information includes six figures and can be found with this article online at http//

## KEY RESOURCES TABLE

| REAGENT or RESOURCE | SOURCE | IDENTIFIER |
| --- | --- | --- |
| **Bacterial strains** |  |  |
| *E. coli* MRE600 | Lab stock | N/A |
| *E. coli* MG1655 | Lab stock | N/A |
| *E. coli* BL21 (DE3) | New England Biolabs | Cat# C2527H |
| *E. coli* BX41 (ΔrbfA) | (Xia et al., 2003) | N/A |
| *E. coli* BW25113 | CGSC, Yale University | CGSC#7636 |
| *E. coli* JW0050-3 (ΔksgA) | (Baba et al., 2006) | CGSC#8292 |
| *E. coli* TPR201 | (Andrésson and Davies, 1980) | CGSC#8265 |
| *E. coli* JK382 | (Sussman et al., 1996) | CGSC#7503 |
| *E. coli* JK378 (*infC362*) | (Sussman et al., 1996) | CGSC#7501 |
| **Plasmids** |  |  |
| pET21b-RbfA-A2C | GenScript | N/A |
| p15BHA | (Dammel and Noller, 1995) | N/A |
| p15B-RbfA-A2C-HA | This study | N/A |
| pGEMX2His-RsgA | (Goto et al., 2011) | N/A |
| pProEx-HTb-IF1 | (Simms et al., 2014) | N/A |
| pET24b-IF2 | (Brunelle et al., 2006) | N/A |
| pProEx-HTb-IF3 | (Simms et al., 2014) | N/A |
| pProEx-HTb-IF3-K110L | This study | N/A |
| pProEx-HTb-IF3-Nter | This study | N/A |
| pProEx-HTb-IF3-Cter | This study | N/A |
| IF3.C65S.S38C.K97C | (Elvekrog and Gonzalez, 2013) | N/A |
| IF3-Y75N.C65S.S38C.K97C | (Elvekrog and Gonzalez, 2013) | N/A |
| **Chemicals/reagents** |  |  |
| 4-20% SDS PAGE | Bio-Rad | Cat# 4561093 |
| acrylamide: bis-acrylamide (37.5:1) | Bio-Rad | Cat #1610148 |
| MOPS buffer | Teknova | Cat #M2101 |
| K<sub>2</sub>HPO<sub>4</sub> | Teknova | Cat #M2102 |
| PVDF membrane | Novex | Cat #LC2002 |
| Cy3-maleimide dye | GE Healthcare | Cat # PA23031 |
| Cy5-maleimide dye | GE Healthcare | Cat # PA25031 |
| S-35-Methionine | Perkin Elmer | Cat #NEG009T001MC |
| 3H-Uridine | Perkin Elmer | Cat #NET174001MC |
| ATP [γ-32P] | Perkin Elmer | Cat #BLU502Z500UC |
| **Critical Commercial Assays** |  |  |
| PURExpress Δribosome kit | New England Biolabs | Cat # E3313S |
| RNeasy kit | Qiagen | Cat # 74104 |
| **Oligonucleotides** |  |  |
| Primer 161 | (Moazed et al., 1986) | N/A |
| Primer 323 | (Moazed et al., 1986) | N/A |
| Primer 1523 | (Seistrup et al., 2017) | N/A |
| Cy3-SA5 | (Kim et al., 2014) | N/A |
| **Antibodies** |  |  |
| Anti-IF3 | (Connolly and Culver, 2013) | N/A |
| goat anti-rabbit WesternDot-655 | ThermoFisher | Cat# W10814 |
| **Software** |  |  |
| ImageQuant | GE Healthcare | N/A |
| Inkscape |  | N/A |

## CONTACT FOR REAGENT AND RESOURCE SHARING

Further information and requests for resources and reagents should be directed to and will be fulfilled by Sarah A. Woodson.

## METHOD DETAILS

### Strains and culture conditions

Strains are listed in the Key Resources Table. All bacterial strains were grown in LB media unless stated otherwise. Media were supplemented with antibiotics (100 µg/ml ampicillin, 25 µg/ml kanamycin, 10 µg/ml tetracycline, and 100 µg/ml kasugamycin) as required.

Growth analysis: *E. coli* strains JK382 (parental) and JK378 (*infC362*) were transformed with either p15BHA over-expressing RbfA (Dammel and Noller, 1995; Sussman et al., 1996) or with empty vector (pSE420), and were grown overnight at 37 °C in LB media supplemented with 10 µg/ml tetracycline plus 100 µg/ml ampicillin. These strains were sub-cultured in fresh media containing 50 µg/ml ampicillin and the optical density (OD_600nm_) was recorded at the indicated intervals. For growth analysis under nutrient limited conditions, cells were grown overnight as above and sub-cultured into minimal media:1X MOPS (Teknova # M2101), 2 mM K_2_HPO_4_ (Teknova #M2102), 0.1 µg/ml thiamine, 0.4% glucose, supplemented with antibiotics as above as previously reported (Mechold et al., 2002).

### Plate assays

For plate assays, a single colony from a freshly prepared plate was re-streaked on plates containing LB agar and antibiotics, and incubated at either 37 °C or 18 °C. Plates were imaged after 1-3 days. For the *rbfA* complementation assay, p15B-RbfA-A2C-HA was transformed into BX41 (*ΔrbfA*) cells. MRE600, BX41 and (BX41*/* p15B-RbfA-A2C-HA) strains were grown on LB agar plates with no antibiotics, 25 µg/ml kanamycin, and 100 µg/ml ampicillin plus 25 µg/ml kanamycin, respectively, in duplicates. Plates were grown at 37 °C or 22°C, and imaged after 1-2 days.

### Plasmids

Plasmids are listed in the Key Resources Table. QuikChange mutagenesis and domain separation was carried out in pProEx-HTb-IF3 to obtain pProEx-HTb-IF3-K110L (primers-IF3-K110L-F 5’AGGTATTACTCCGCAGCCTGATTC3’ and IF3-K110L-R 5’GGAGTAATACCTGATAGTCGCCTTC3’) and pProEx-HTb-IF3-Nter (primers-IF3-Nter-F 5’TAACTCGAGGCATGCGG3’ and IF3-Nter-R 5’GATAACTTTTTGCTTTTTCTTCTG3’) and pProEx-HTb-IF3-Cter (primers-IF3-Cter-F 5’CAGGTTAAGGAAATTAAATTCCG3’ and IF3-Cter-R 5’AATGGATCCCATGGCG3’). QuikChange mutagenesis was also carried out on plasmid p15BHA to obtain p15B-RbfA-A2C-HA (primers-p15BHA-F 5’GAATAAACCATGGATGTGCAAAGAATTTGGTCGCC3’ and p15BHA-R 5’GGCGACCAAATTCTTTGCACATCCATGGTTTATTC3’) for the RbfA complementation assay.

### Ribosome purification

Mature ribosomes were purified from MRE600 following the protocol of (Spedding, 1990). Near mature 30S (nm30S) subunits were isolated from JW0050-3 *(ΔksgA)* and TPR201 cells following the same protocol, except that the cell lysis was performed in an ethanol-dry ice bath. Pre-30S subunits were purified as follows: BX41 (*ΔrbfA*) cells were grown in 5 ml LB at 37 °C for 9 h with shaking at 250 rpm. A 2 ml aliquot was transferred to 200 ml LB supplemented with 25 µg/ml kanamycin and grown overnight at 37 °C with shaking at 250 rpm. 50 ml of this culture was then transferred to two flasks each containing 2 l LB supplemented with 25 µg/ml kanamycin. The cultures were grown at 37 °C until O.D._600_ ∼0.7, after which the cultures were shifted to a shaking incubator pre-cooled at 17 °C and grown until O.D._600_ ∼ 1.2 (ClatterbuckSoper et al., 2013). The cultures were stored at 4 °C for 30 min before pelleting cells at 4,000 × *g* for 10 min at 4 °C. The cell pellets were washed once with 10 mM Tris-HCl pH 7.5, 15 mM MgCl_2_ and re-pelleted as above and stored at −20 °C until needed. The pellets were resuspended in 15 ml lysis buffer containing 10 mM Tris-HCl pH 7.5, 15 mM MgCl_2_ and 1 mg/ml lysozyme (hen egg white; Sigma #L7651) (freshly prepared). The cells were incubated for 5 min with intermittent vortexing, frozen in an ethanol-dry ice bath, and allowed to thaw at room temperature. Cell lysates were cleared twice by centrifugation at 20,000 × *g* for 20 min at 4 °C. 10-40% sucrose gradients in 60 mM Tris-HCl pH 7.5, 30 mM MgCl_2_, 300 mM NH_4_Cl, 6 mM DTT, 3 mM β-mercaptoethanol were prepared in SW28 rotor tubes (Beckman Coulter) using a gradient master (BioComp). An equal quantity of cleared lysate (OD_260nm_ ∼100) was layered onto each gradient and spun at 25,000 rpm for 2 hours at 4 °C (SW28 rotor; Beckman Coulter). The fraction corresponding to pre-30S particles was collected using a fractionator (BioComp) and analyzed for the presence of 17S rRNA on a 1.5% agarose-TAE gel. The protein component was analyzed by 4-20% SDS-PAGE (Bio-Rad #4561093). Pooled fractions were concentrated using 100 kDa MWCO ultrafiltration tubes (Millipore # UFC510096) and exchanged 3-5 times with buffer A (20 mM Tris-HCl pH7.5, 40 mM NH_4_Cl, 60 mM KCl, 10 mM MgCl_2_). Aliquots were snap frozen in liquid nitrogen and stored at −80 °C until further use.

### Protein purification and fluorescence labeling

*E. coli* proteins S1, RbfA, RsgA, IF1, IF2, and IF3 (WT, mutant and domains) with histidine tags were purified from BL21(DE3) cells harboring the respective plasmids listed in the Key Resources Table. Over-expression was induced with 1 mM IPTG at 37 °C. Pellets were lysed in lysis buffer (50 mM Tris-HCl pH7.5, 500 mM NaCl) using sonication, and cell debris was removed by centrifugation at 20,000 × *g* for 20 min. Cleared lysates were passed through a 5 ml HiTrap Ni-column (GE Healthcare #17040901) pre-equilibrated with lysis buffer. The column was washed with 25 column volumes of lysis buffer containing 30 mM imidazole and 25 column volumes of lysis buffer with 30 mM imidazole plus 1 M NaCl. His-tagged protein was eluted using lysis buffer plus 500 mM imidazole. Protein-containing fractions were pooled and dialyzed in 50 mM Tris-HCl pH 7.5, 300 mM KCl, 10 mM MgCl_2_, 6 mM β-mercaptoethanol, 10% glycerol after the purity was checked by SDS PAGE. For fluorophore labeling, the protein of interest containing a single cysteine was dialyzed overnight at 4 °C against 80 mM Tris-HCl pH 7.5, 1 M KCl, 1 mM TCEP. A maleimide conjugate of Cy5 or Cy3 dye (GE Healthcare#PA25031, PA23031) was dissolved in anhydrous DMSO (Molecular Probes #D12345), mixed with the protein of interest following the manufacturer’s instruction and incubated for 2-3 h at room temperature in the dark. Reactions were quenched with 6 mM β –mercaptoethanol, and the labeled protein was re-purified through Ni-column, concentrated in 3 kDa MWCO filtration units (Millipore#UFC900324) and exchanged with dialysis buffer as described above. Subsequently, concentrated proteins were dialyzed a second time against a large excess of dialysis buffer, aliquoted, snap frozen and stored at –80 °C. The labeling efficiency and concentration were checked by UV absorption (Nanodrop; Thermo Scientific). IF3 was used before and after removing the histidine tag by TEV protease.

### Ultracentrifugation pelleting assay and Ultrafiltration assay

200 nM pre-30S or 30S subunits was incubated with 400 nM Cy5-RbfA in buffer A with 3 mM DTT (Buffer A*) for 15 min at 37 °C (300 µL total volume is sufficient for 6 pelleting assays). Unbound Cy5-RbfA was removed by passing the reaction mixture through a 100 kDa MWCO filtration unit (Millipore#UFC510096) for 5 min at 10,000 x *g*. The filter was washed twice with 200-400 µl buffer A* by centrifugation as above and the complexes were recovered by centrifugation at 1,000 x *g* following the manufacturer’s instructions. To monitor the release of Cy5-RbfA in a pelleting assay (Jeganathan et al., 2015; Shoemaker and Green, 2011), the pre-30S or 30S•Cy5-RbfA complex (50 µL) was incubated with excess IF1/IF2/IF3/mRNA/S1 (individually or in combination) in buffer A* for 15 min at 37 °C in a 100 µl reaction volume. 10 µl of this mixture was saved as “input” and 90 µl was separated through a 2 ml 1.1 M sucrose cushion in buffer A by ultracentrifugation in a SW50 rotor (Beckman Coulter). The pellet was dissolved in 30 µl buffer A*, and 10 µl of the input and pellet samples was mixed with 2 µl 2X Tricine loading dye (Bio-Rad#161-0739) and boiled for 2 min at 95 °C and resolved by 4-20% SDS PAGE. Gels were scanned for Cy5 signal using a Typhoon imager (GE Healthcare) and quantified using ImageQuant (GE Healthcare) before staining with Coomassie blue. For monitoring release with a filtration assay as previously described (Goto et al., 2011; Jeganathan et al., 2015), unbound Cy5-RbfA was removed from the release reaction mixture by passing the reaction mixtures again through a 100 kDa MWCO filtration unit as above, and input and retentate was further processed similar to pelleting assay.

30S assembly intermediates were prepared by mixing 100 nM 17S rRNA (transcribed) with purified recombinant primary assembly r-proteins (S4, S7, S8, S15, S17, and S20) or primary plus secondary r-proteins (S4, S7, S8, S15, S17, and S20 plus S6, S16, S9, S13, S18, and S19) (200 nM) as previously described (Culver and Noller, 1999; Traub and Nomura, 2006) in HKM20 buffer (80 mM Tris-HCl pH 7.8, 330 mM KCl, and 20 mM MgCl_2_) at 42 °C for 1 h. 17S rRNA or native 30S subunits was also incubated with Cy5-RbfA separately as controls. The reaction mixtures were then layered onto 1.1 M sucrose cushions and analyzed as described above.

### Primer extension

Primer extension on total RNA extracted from *E. coli* (BW25113) and BX41 (Δ*rbfA*) or rRNA purified from 30S fractions was performed using primers 5′ labeled with either ^32^P-or Cy5. The 16S 5′ end was analyzed with primer 161 (5’GCGGTATTAGCTACCGT3’) or primer 323 (5’ AGTCTGGACCGTGTCTC3’) as described previously (ClatterbuckSoper et al., 2013). For checking the methylation of 16S A1518 and A1519, 5′-^32^P-labeled primer 1523 (5’GGAGGTGATCCAACCGC3’) (Seistrup et al., 2017) was used on rRNA purified from immature and mature 30S subunits. In all cases, the rRNA was purified using RNeasy kit (Qiagen# 74104).

### *In vitro* translation assay

Purified pre-30S, nm30S or 30S subunits and native 50S subunits were incubated with or without 3 µM RbfA and combined with *in vitro* translation components (PURExpress; New England Biolabs # E3313S). Each reaction mixture (15-20 µl) was supplemented with 1 µl ^35^S methionine (1 µCi, Perkin Elmer #NEG009T001MC) and 1 µl RNase inhibitor (Promega#N2615) and incubated at 37 °C for 2 h in an incubator. The reaction was stopped by placing tubes on ice for 5 min, followed by the addition of 2 µl 2X Tricine loading dye (Bio-Rad#161-0739). 8-10 µl of each reaction was loaded without boiling onto a 4-20% SDS PAGE. The gel was then dried under vacuum, exposed overnight against a phosphor screen and imaged using a Typhoon scanner.

### ^3^H-Uridine pulse labeling and polysome profiling

BW25113 cells were first grown overnight in LB at 37 °C with shaking at 250 rpm, then transferred to fresh 5 ml LB at 37 °C with shaking at 250 rpm. After 3 h or after 10 h, 50 µl ^3^H-Uridine (50 µCi, PerkinElmer#NET174001MC) was added to the culture, which was allowed to grow for another 9 min, before the addition of 25 µg/ml chloramphenicol for 1 min. Cells were harvested, washed with cold 10 mM Tris-HCl pH7.5 and 15 mM MgCl_2_, and pellets stored at –80 °C. To analyze polysome profiles, cell lysate was prepared as above. Equal amounts of cleared lysate containing total ribosomes (OD_260nm_ ∼20) was separated through 10-40% sucrose gradient in the buffer as above for 2.5 h at 35,000 rpm in a SW41 rotor (Beckman Coulter). The gradients were analyzed using a fractionator (Biocomp) and the A_260nm_ recorded. Fractions (250 µl) covering the entire gradient were collected in 96-well plate and ^3^H-uridine incorporation was measured by liquid scintillation counting (Beckman). rRNA analysis was performed using primer extension as described above.

### Western blot

Western blot analysis was performed as previously described (Connolly and Culver, 2013; Goto et al., 2011). Briefly, reaction mixtures (0.5-1 µg), cell lysates (40-60 µg) or 30S fractions (5-10 µg) (concentrated using 3 kDa MWCO filtration tubes) from polysome profiling experiments were separated by 4-20% SDS PAGE and transferred to a PVDF membrane (Novex#LC2002) and probed with polyclonal antibodies against IF3. Antibody binding was detected using a secondary goat anti-rabbit WesternDot-655 antibody (ThermoFisher, W10814), following the manufacturer’s protocol. Membranes were imaged using a Typhoon scanner.

### Native PAGE experiments

Co-localization assays were performed by incubating Cy5-RbfA and Cy3-IF3, together or separately, with or without pre-30S (*ΔrbfA*), 30S (MRE600), nm30S (*ΔksgA*) or nm30S (TPR201) subunits (100 nM each reactant) in buffer A* for 15 min at 37 °C in a 10 µl reaction mixture. 2 µl native loading dye (50% sucrose, 0.02% bromophenol blue) was added before loading 8 µl on a native 4% PAGE (acrylamide: bis-acrylamide 37.5:1, Bio-Rad#1610148). The gel was run at ∼15.5 °C for ∼45 min in 1X THEM buffer (56 mM Hepes-K^+^, 34 mM Tris-HCl pH 7.5, 5 mM MgCl_2_, 0.1 mM EDTA) before scanning with 532 nm excitation (Cy3) and an emission filter at 610 nm for FRET (Typhoon). The FRET efficiency (E) was calculated from (Sabanayagam et al., 2004).

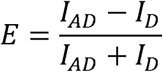

where *I*_*AD*_ is the intensity of the acceptor in the presence of the donor (i.e. FRET) and *I*_*D*_ is the intensity of the donor. The gels were also imaged with 633 nm excitation (Cy5).

For binding of RbfA with the 16S rRNA 5’ domain, an extended 5’domain RNA was transcribed and annealed with Cy3-SA5 oligonucleotide as previously described (Abeysirigunawardena et al., 2017; Kim et al., 2014). The 5′ domain•Cy3-SA5 complex was incubated with Cy5-RbfA, with or without 5’ domain binding ribosomal proteins (S4, S17, S20, and S16), at 37 °C for 15 min in HKM20 buffer. For measuring binding of RbfA with 16S (native) or 17S rRNA (transcribed), increasing concentrations of 16S (10-100 nM) or 17S (20-100 nM) rRNA was incubated with 100 nM Cy5-RbfA at 37 °C for 15 min in HKM20 buffer. The complexes were resolved and analyzed as described above.

For binding of RbfA with the 30S translation initiation complex (30SIC), 30S subunits were first incubated at 42 °C for 30 min in an initiation complex buffer (ICB) containing 50 mM Tris-HCl pH 7.5, 70 mM NH_4_Cl, 30 mM KCl, 7 mM MgCl_2_ as previously described (Julián et al., 2011). The initiation complex was formed by combining 100 nM 30S subunits, 500 nM mRNA (*sodB*), 300 nM fMet-tRNA^fmet^, 300 nM each IF1, IF2 and IF3, and 300 nM GTP in ICB at 37 °C for 15 min. Subsequently, 100 nM Cy5-RbfA was incubated with 50 nM initiation complex in ICB at 37 °C for 15 min. 2 µl native loading dye was added and the reaction mixtures were resolved and analyzed as described above.

