## Supporting Information for "IF3 licenses newly made 30S subunits for translation during stress"

Indra Mani Sharma and Sarah A. Woodson*
T. C. Jenkins Department of Biophysics, Johns Hopkins University, 3400 N. Charles St., Baltimore, MD 21218, USA

Six (6) figures associated with main figures and discussion.

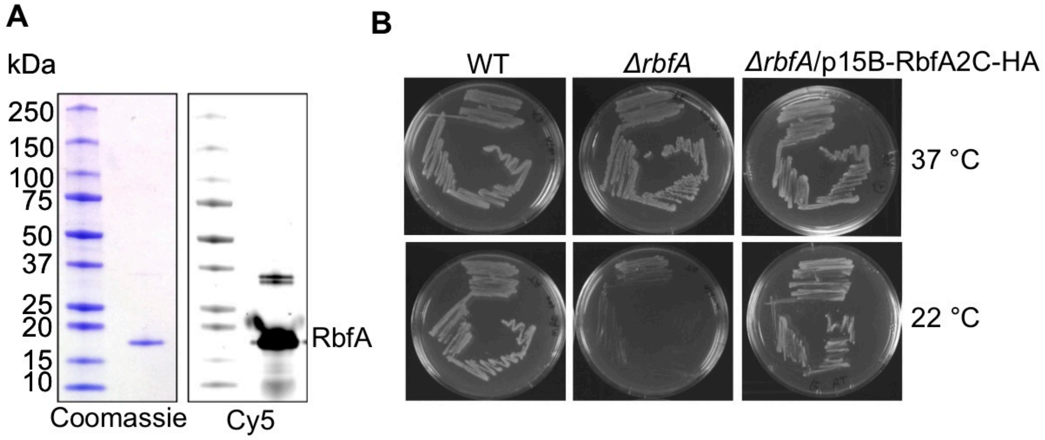


**Figure S1 (related to Figure 1) Preparation of Cy5-labeled RbfA.**
(A) 4-20% SDS PAGE showing the purity of RbfA:A2C protein after labeling with Cy5-maleimide.
(B) The *in vivo* function of RbfA:A2C was tested by complementation of the cold sensitive phenotype of BX41 *ΔrbfA* strain by expression of the *rbfA*:A2C mutant from plasmid p15B-RbfA2C-HA. The mutant RbfA was able to restore growth at 22 ˚C, comparable to an *rbfA*$^{+}$ *E.coli* strain (MRE600).

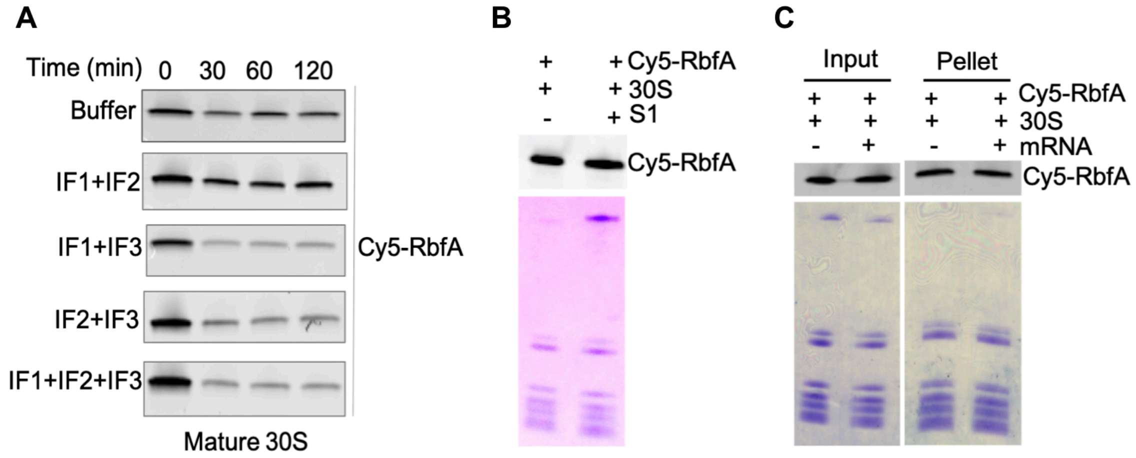


**Figure S2 (related to Figure 1) Other 30SIC components do not release RbfA.**
(A) Ultrafiltration assays showing the kinetics of release of Cy5-RbfA from mature 30S subunits in the presence of buffer or combinations of IF1, IF2 and IF3. The retentates were analyzed by SDS-PAGE and scanned for Cy5 intensity as described in the Methods (also see Figure 1). The “0” min sample was filtered once to remove excess free Cy5-RbfA at the start of the assay. In the following lanes, the sample was filtered again at 30, 60 and 120 min to remove any RbfA that dissociated during this period. Top, the RbfA•30S complex is stable for 2 h in the absence of release factors. Lower panels, the presence of IF1 and IF2 had no effect on the release activity of IF3.
(B) An ultrafiltration assay showing that addition of excess ribosomal protein bS1 (4 μM) cannot displace Cy5-RbfA from 30S subunits. The retentate was resolved on a 4-20% SDS-PAGE and scanned for Cy5 intensity (top) or stained with Coomassie (bottom).
(C) A pelleting assay showing that *sodB* mRNA (0.4 μM), which contains a strong Shine-Dalgarno sequence, cannot displace Cy5-RbfA from 30S subunits.

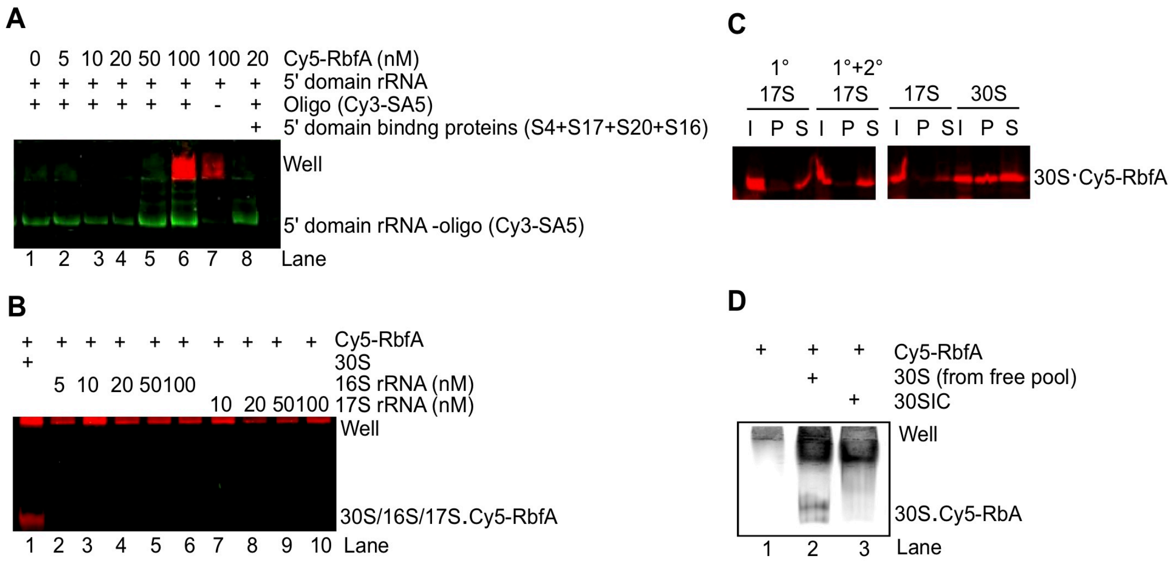


**Figure S3 (related to Figure 3) Early 30S assembly complexes are not recognized by RbfA.**

(A) Native 4% PAGE showing that Cy5-RbfA cannot bind with the 5' domain of 16S rRNA. 5' domain RNA (15 nM) with a 3' extension was hybridized with a complementary Cy3-SA5 DNA oligomer (10 nM) as previously described (Abeysirigunawardena and Woodson, 2015; Kim et al., 2014). 5' domain RNA•Cy3-SA5 complexes were incubated without or with an increasing concentration of Cy5-RbfA. Lane 8, reaction includes 2.5 μM each 5' domain proteins S4, S17, S20, and S16 (Abeysirigunawardena et al., 2017) to see if they could promote RbfA binding with 5' domain RNA.

(B) Native 4% PAGE showing that Cy5-RbfA cannot bind with 16S rRNA (native) and 17S rRNA (transcribed). Lane 1, 30S subunits were used as a control, in which free Cy5-RbfA remains in the well and bound Cy5-RbfA migrates into the gel together with 30S complexes.

(C) A pelleting assay showing that Cy5-RbfA cannot bind transcribed 17S rRNA reconstituted with primary (1˚) or primary and secondary (1˚+ 2˚) 30S proteins (Culver and Noller, 1999).

(D) Native 4% PAGE showing that Cy5-RbfA binds free 30S subunits (lane 2) but not translation initiation complexes (30SIC) (lane 3). 30SIC was formed by incubating 30S subunits with initiation factors, mRNA (*sodB*), and fMet-tRNA$^{fmet}$ as previously described (Julián et al., 2011).

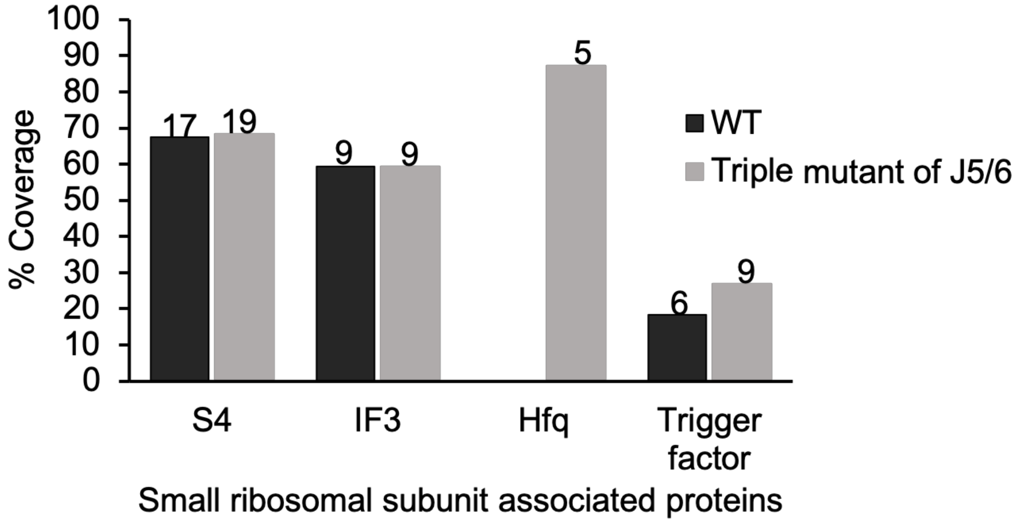


**Figure S4 (related to Figure 3) Detection of IF3 in a pre-30S complex.**
Qualitative mass spectrometry data showing co-purification of IF3 with 30S (black bars) and OFF-pathway MS2-tagged pre-30S particles (gray bars). These pre-30S subunits have three mutations (A59, 60C, G107C) in helix junction J5/6 of the 16S rRNA. Data reanalyzed from (Sharma et al., 2018). The number of peptides observed for each protein are shown on the top of respective bar.

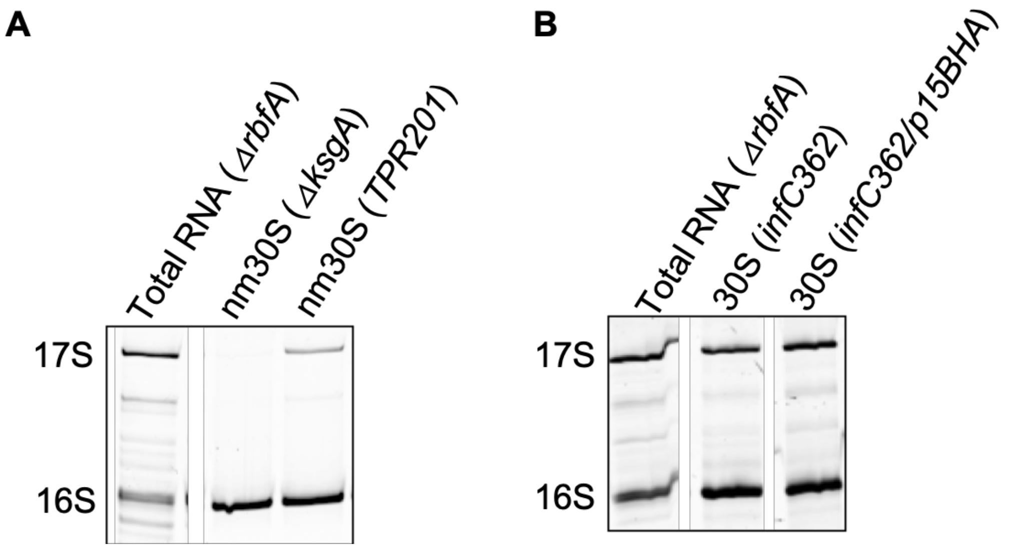


**Figure S5 (related to Figure 4 and 5) Processing of 16S rRNA in mutant strains.** Processing of the 16S 5′ end was analyzed by primer extension (Cy5-labeled primer 161) and denaturing 8% PAGE. The analysis was done with total cellular RNA or 16S rRNA from 30S complexes.

(A) The 16S rRNA from nm30S subunits was fully processed in Δ*ksgA* and ≥90% processed in TPR201 cells.

(B) There is similar proportion of 17S and 16S rRNA in 30S fractions from *infC362* and *infC362*/p15BHA strains. Polysome profiles were analyzed from cells grown in rich LB media until $OD_{600nm}$ ~0.2.

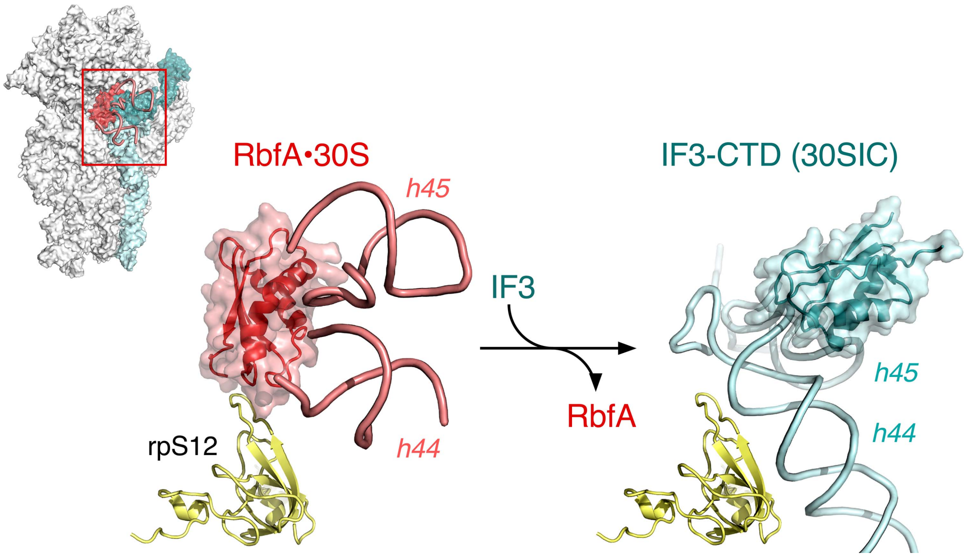


**Figure S6 (related to discussion) RbfA and IF3 modulate the conformation of 16S H44 and H45.**

Binding of RbfA (red) unfolds the top of 16S H44 and H45 (salmon) in the RbfA•30S complex (Datta et al., 2007). By contrast, binding of IF3 (teal) and extension of its CTD stabilizes the docked conformation of 16S H44 and H45 (cyan) in 30S initiation and pre-initiation complexes (Hussain et al., 2016; López-Alonso et al., 2017). The opposing actions of RbfA and the IF3 CTD may explain how IF3 can promote the release of RbfA when H44 and H45 are correctly folded. Ribosomal protein uS12 (yellow) is shown for reference. Inset upper left: RbfA and 16S fragments modeled into the cryo-EM density map of the 30S•RbfA complex (PDB: 2R1G; (Datta et al., 2007)) were superimposed on coordinates from a cryo-EM model of 30SIC2 (PDB: 5ME1; (López-Alonso et al., 2017)). Atoms in 16S H1, H18, and H27 were aligned using PyMOL. Ribbons for IF1, IF2 and fMet-tRNA$^{init}$ in the 30IC2 are not shown for clarity.
